# Brain cell-specific origin of circulating microRNA biomarkers in experimental temporal lobe epilepsy

**DOI:** 10.1101/2023.06.02.542426

**Authors:** Elizabeth Brindley, Mona Heiland, Catherine Mooney, Mairead Diviney, Omar Mamad, Thomas D. M. Hill, Yan Yan, Morten T. Venø, Cristina Ruedell Reschke, Aasia Batool, Elena Langa, Amaya Sanz-Rodriguez, Janosch Heller, Gareth Morris, Karen Conboy, Jørgen Kjems, Gary P. Brennan, David C. Henshall

## Abstract

The diagnosis of epilepsy is complex and challenging and would benefit from the availability of molecular biomarkers, ideally measurable in a biofluid such as blood. Experimental and human epilepsy are associated with altered brain and blood levels of various microRNAs (miRNAs). Evidence is lacking, however, as to whether any of the circulating pool of miRNAs originates from the brain. To explore the link between circulating miRNAs and the pathophysiology of epilepsy, we first sequenced argonaute 2 (Ago2)-bound miRNAs in plasma samples collected from mice subject to status epilepticus induced by intraamygdala microinjection of kainic acid. This identified time-dependent changes in plasma levels of miRNAs with known neuronal and microglial-cell origins. To explore whether the circulating miRNAs had originated from the brain, we generated mice expressing FLAG-Ago2 in neurons or microglia using tamoxifen-inducible *Thy1* or *Cx3cr1* promoters, respectively. FLAG immunoprecipitates from the plasma of these mice after seizures contained miRNAs, including let-7i-5p and miR-19b-3p. Taken together, these studies confirm that a portion of the circulating pool of miRNAs in experimental epilepsy originates from the brain, increasing support for miRNAs as mechanistic biomarkers of epilepsy.

## Introduction

Epilepsy is a chronic neurological disease characterised by recurring seizures that affects more than 50 million people worldwide (Moshe et al., 2015). Temporal lobe epilepsy (TLE) is the most common and intractable form of the disease in adults. Diagnosis of epilepsy is principally based on clinical history and examination, supported by electroencephalogram (EEG), and other investigational tools such as brain imaging. This requires specialist training, is time-consuming, error-prone and expensive (Simonato et al., 2021). The diagnostic yield from genome sequencing is increasing for many epilepsy syndromes, but is likely to remain low in the TLE population (Kearney et al., 2019). Rates of misdiagnosis remain substantial (Ferrie, 2006;Oto, 2017;Gasparini et al., 2019), with psychogenic non-epileptic seizures (Gasparini et al., 2019) and syncope (Zaidi et al., 2000), adding to the challenge of identifying epilepsy as the underlying cause. Additional tools to support early and accurate diagnosis would reduce the healthcare and socioeconomic burden of epilepsy and improve the quality of life of patients (Ferrie, 2006;Moshe et al., 2015;Oto, 2017;Kearney et al., 2019).

The identification of molecule(s) within an easy-to-access biofluid such as blood would revolutionize the diagnosis of epilepsy (Engel et al., 2013). To be practical, however, such molecules must be stable, simple and cheap to measure, ideally in a point-of-care setting, and be mechanistically linked to the pathophysiology (Enright et al., 2018). For this reason, microRNAs (miRNAs) are promising circulating biomarkers to support epilepsy diagnosis. MiRNA are short noncoding RNAs that post-transcriptionally regulate the gene expression landscape. To function, mature miRNAs are loaded into a binding pocket of an argonaute (AGO) protein, of which AGO2 is a key effector, thereby forming an RNA-induced silencing complex (RISC) (Peters and Meister, 2007). The miRNA-loaded RISC searches for seed regions of nucleotide complementarity along target mRNAs (Chandradoss et al., 2015) and promotes translation inhibition or degradation of the target (Bartel, 2018). Extracellular AGO-bound miRNAs are present within biofluids including plasma (Arroyo et al., 2011;Turchinovich et al., 2011), probably released from diverse tissues (Turchinovich and Burwinkel, 2012), and their altered composition may have diagnostic value (Geekiyanage et al., 2020).

The brain expresses the greatest diversity of miRNAs, with several unique to the brain and to specific brain cell types (Bak et al., 2008;Jovicic et al., 2013;Nowakowski et al., 2018). This contributes to the establishment and functional properties of neuronal networks (Soutschek and Schratt, 2023). There is extensive dysregulation of the miRNA system within structures such as the hippocampus in experimental and human epilepsy (Brennan and Henshall, 2020), and functional studies show targeting certain miRNA has therapeutic potential (Morris et al., 2021). Circulating miRNA levels are altered in epilepsy (Enright et al., 2018). Plasma from rodents subject to evoked seizures and animals with chronic epilepsy display elevations in brain-enriched and inflammation-associated miRNAs (Gorter et al., 2014;Roncon et al., 2015;Brennan et al., 2018). Several of the same miRNAs are differentially expressed in baseline or post-seizure blood samples from patients with drug-resistant epilepsy (Brennan et al., 2018;Enright et al., 2018;Leontariti et al., 2020;Martins-Ferreira et al., 2020), including in AGO-bound fractions (Raoof et al., 2018). It remains unproven, however, whether any of the circulating pool of miRNAs actually originated in the brain. Resolving this issue is a priority to ensure mechanistic links to the underlying pathophysiology.

Here, we report time-dependent changes in brain- and brain-cell type-enriched Ago2-bound miRNAs in the plasma of mice following seizures. Using transgenic mice expressing FLAG-tagged Ago2 under promoters to restrict expression to brain cell types, we report a portion of circulating miRNAs originates from the brain.

## Results

### Circulating Ago2-bound miRNAs in mouse plasma

AGO-bound miRNAs represent a major pool of circulating miRNAs (Arroyo et al., 2011), and measuring AGO-bound miRNAs may enhance the ability to distinguish brain disease subtypes (Raoof et al., 2017), and post-seizure from baseline samples in plasma from epilepsy patients (Raoof et al., 2018). Moreover, there is disruption of the blood-brain barrier after seizures sufficient to allow passage of macromolecules between the blood and brain (van Vliet et al., 2007;Reschke et al., 2021).

To obtain evidence that seizures and the pathophysiologic changes that accompany the development of epilepsy are associated with changes to the circulating AGO pool of miRNAs, we first sequenced Ago2-bound miRNAs from the plasma of mice following status epilepticus (SE) triggered by intraamygdala microinjection of kainic acid (Mouri et al., 2008;Brennan et al., 2018). The model displays typical features of the pathophysiology of TLE including select neuron loss and gliosis within the hippocampus and the emergence of recurrent, spontaneous seizures which are resistant to various anti-seizure medicines (Mouri et al., 2008;West et al., 2022). Figure 1A shows a schematic of the study design, including time points at which blood was collected. In order to obtain sufficient Ago2 yield for sequencing, 500 µl of plasma was used, from pooling samples from n = 10 mice per time point. After confirming immunoprecipitation of Ago2 from mouse plasma (Supplementary Figure S1A), plasma samples at various time points were processed for Ago2 elution followed by small RNA sequencing on an Illumina HiSeq.

**Figure 1.**
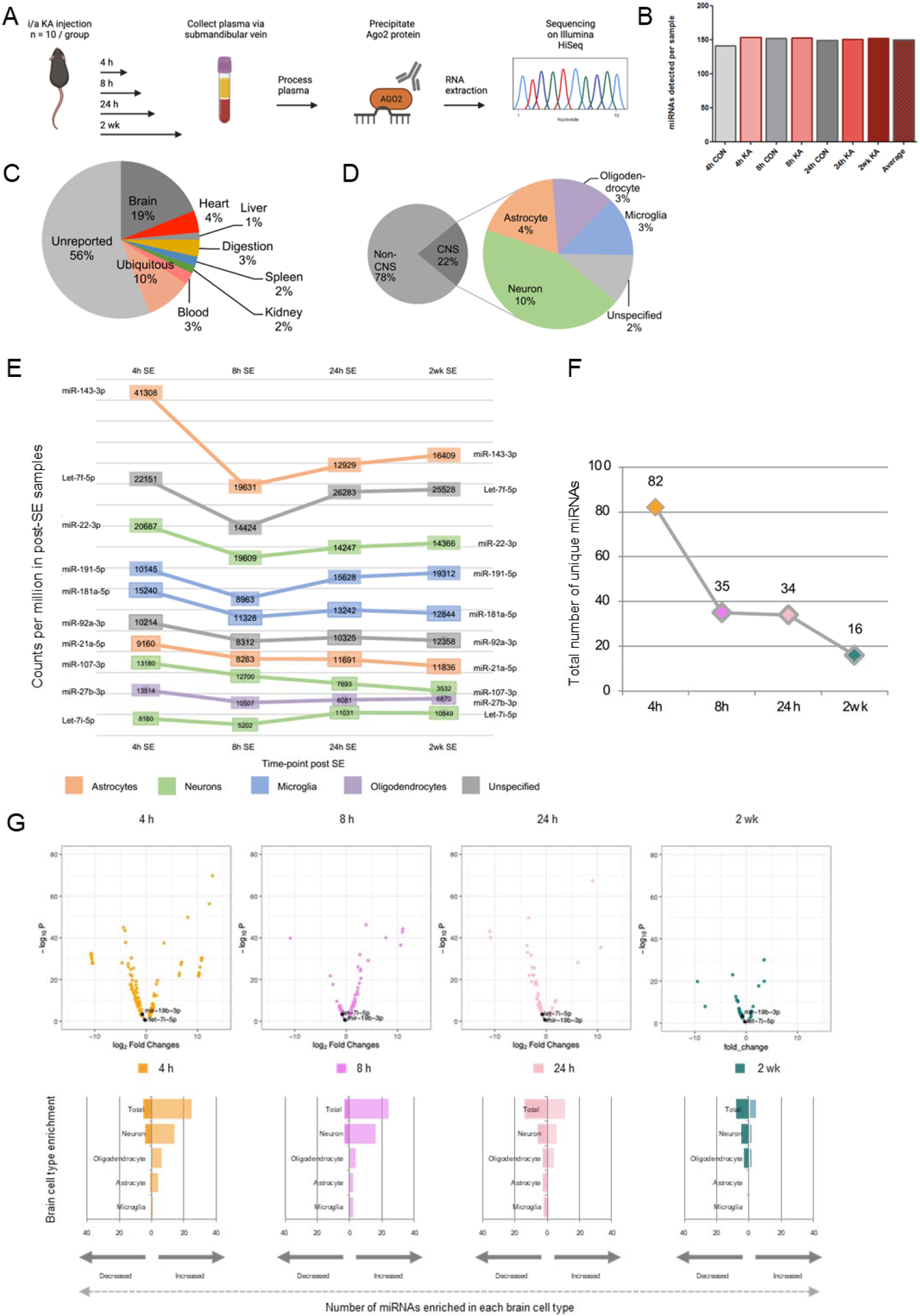
Temporal and brain origins of circulating Ago2-bound miRNAs following SE in mice. (A) Schematic of the workflow for generating samples and small RNA sequencing on circulating Ago2-bound miRNA across time-points following SE in mice. (B) Total number of unique miRNAs present in all time points (4 h, 8 h, 24 h and 2 weeks post-treatment. (C) Percentages of reported tissue enrichment of miRNA present in the Ago2 fraction of mouse plasma across all time points. (D) Percentage of reported non-CNS, CNS and specific brain cell type-(neuron-, microglia-, astrocyte, oligodendrocyte-) enrichment of miRNA present in the Ago2 fraction of mouse plasma (control + SE) across all time points (4 h, 8 h, 24 h, 2 weeks). (E) Counts per million of top 10 most abundant brain cell type enriched miRNA observed in SE across all time points. (F) Numbers of differentially expressed miRNA in the Ago2 fraction of mouse plasma across time-points. (G) Log_2_ fold change differences between control and SE at 4 h, 8 h, 24 h and 2 weeks plotted against –log_10_ *p*-values for all miRNAs detected in Ago2 fraction of mouse plasma. miRNAs of particular interest are labelled.

A total of 23.7 million reads were mapped from 7 samples across four time points in control and SE mice. The numbers of normalised miRNA counts per sample varied from 2,038,033 to 5,388,335, with an average of 3,390,383 (Supplementary Figure S1B). Numbers of unique miRNA detected per sample showed minimal variation (range 141-154; average: 150) (Figure 1B). Likewise, the average number of miRNA was quite similar between seizure compared to control mice (Figure 1B). The highest number of counts was observed in the 4 h post SE time-point (5,388,335) (Supplementary Figure S1B). While the lack of replicates prevents a statistical analysis, miRNA counts at the 8 h and 24 h post SE time-points were at intermediate levels (3,151,945 and 3,709,772, respectively) and the lowest numbers of counts (2,532,271) in a sample were observed at the 2-week post SE time-point (Figure 1C). These findings suggest that SE may have a small effect to increase the amount and diversity of circulating Ago2-bound miRNAs, which subsides over time.

### Tissue and cell origins of circulating Ago2-bound miRNAs after SE in mice

To explore the likely tissue origins and diversity (i.e., the number of different miRNAs) of the circulating Ago2-bound miRNAs, we compared the miRNAs across all samples (SE and control) to tissue-enriched miRNA datasets (Lagos-Quintana et al., 2002;Landgraf et al., 2007;Liang et al., 2007;Lee et al., 2008;Bargaje et al., 2010;Guo et al., 2014;Ludwig et al., 2016). Nearly half of the detected miRNAs were predicted to be enriched in a specific tissue type (Figure 1C). MiRNAs enriched in a number of different tissue types included heart (4%), liver (1%), digestive system (3%), spleen (2%), kidney (2%) and blood (3%). The highest proportion of circulating miRNA for which specific tissue expression was known were brain-enriched miRNA (19%).

Next, we used available data on brain cell-enriched miRNAs across all samples (SE and control) to assign the detected miRNAs to cell types, including neurons and glia (*see* Methods and Supplementary Table S1; Figure 1D). Where there were discrepancies in assigning brain cell types, priority of assignment was based on a ranking (*see* Methods). For miRNAs reported as brain-enriched but where there was insufficient information on cell type, these were termed unspecific brain enriched (Ludwig et al., 2016). Of all miRNAs detected in the Ago2 fraction of mouse plasma, 22% had known brain or brain cell type enrichment (Figure 1D). MiRNAs of neuronal origin were the most abundant in the Ago2-eluted plasma of mice, followed by astrocytic, oligodendrocytic and microglial. Numbers of miRNA enriched in these brain cell types are as follows; unspecific (15), neuron (62), astrocyte (26), oligodendrocyte (19) and microglia (18) (Figure 1D). Counts per million of the 10 most abundant brain cell type enriched miRNA observed in SE across all time points include miRNAs with reported enrichment in a variety of brain cell types; astrocytes, neurons, microglial and oligodendrocytes (Figure 1E).

### Evoked seizures in mice alter circulating Ago2-bound miRNAs

We next explored time-dependent changes to the Ago2-bound miRNAs in mouse plasma after SE (Figure 1E, F, G and Supplementary Figure S2, S3). The largest differences were observed at early time-points. At the 4 h, 8 h and 24 h time-points after SE, changes were observed for miRNA enriched in brain, heart, liver, digestive system, spleen, kidney and the blood (Supplementary Figure S2, S3). At the two-week time-point, when mice typically have spontaneous seizures, miRNA enriched in brain, heart, spleen, kidney, and the blood were dysregulated. In addition to these organ-enriched miRNA, miRNA with broad expression across organs were also dysregulated (See Supplementary Table S1).

The largest differences between SE and control samples for Ago2-bound miRNA were found at the earliest time point (4 h; 82 miRNAs) and decreased over time (Figure 1F). Differences were also found at later time points, although numbers declined (35, 34 and 16 miRNAs at the 8 h, 24 h and 2-week time-points, respectively). At the 4 h time-point, 22 and 5 brain/brain cell type-enriched miRNA were higher or lower, respectively, in SE samples compared to control (Figure 1G). Among those showing higher levels were 7 neuron-enriched (e.g., miR-132-5p), 5 astrocyte-enriched (e.g., miR-31-5p) and 1 microglia-enriched (miR-19a-3p). Neuron-enriched miRNAs, including miR-139-5p and miR-135-5p were also among those at lower levels 4 h after SE. At the 8 h time-point, 19 and 3 brain-enriched miRNAs were higher or lower, respectively (Figure 1G). The proportion of the elevated miRNAs that were of likely neuronal origin all increased (e.g., miR-132-3p and miR-218-5p). Brain-enriched miRNAs for which cell-type-enrichment was uncertain were also detected at increased levels compared to controls. By 24 h, a more even mix of higher and lower miRNA levels was observed (Figure 1G). A high proportion of those at higher levels were neuronal (e.g., miR-124-3p and miR-434-3p) whereas several glial-enriched miRNAs were among those at lower levels (e.g., miR-31-3p and miR-29b-3p). At the two-week time-point, a similar ratio of higher and lower miRNAs were observed between SE and control. Again, a majority of those increased being of likely neuronal origin (e.g., miR-669a-5p and miR-541-5p) (Figure 1G). These findings suggest SE adjusts the abundance and cell type-contribution of the circulating pool of Ago2-bound miRNAs in a time-dependent manner in the plasma of mice.

### Neuron- and microglia-restricted epitope-tagged Ago2 mice

To determine if some of the circulating miRNA biomarkers of epilepsy originate from the brain, and to establish whether any of these are from neurons, we generated mice that express a FLAG-tagged Ago2 protein under a neuron or microglial-specific promoter. To achieve this, mice expressing a FLAG-tagged Ago2 with a loxP flanked stop codon under the control of the ubiquitously expressed ROSA26 promoter (Schaefer et al., 2010;Tan et al., 2013), were crossed with transgenic lines with a tamoxifen-inducible cre-recombinase driven by a neuronal promoter, Thy1 (Young et al., 2008), or the microglial Cx3cr1 promoter (Parkhurst et al., 2013) (Figure 2A). Attempts to generate an astrocyte-specific FLAG-Ago2 line using a cre-*Gfap* promoter were unsuccessful (data not shown).

**Figure 2.**
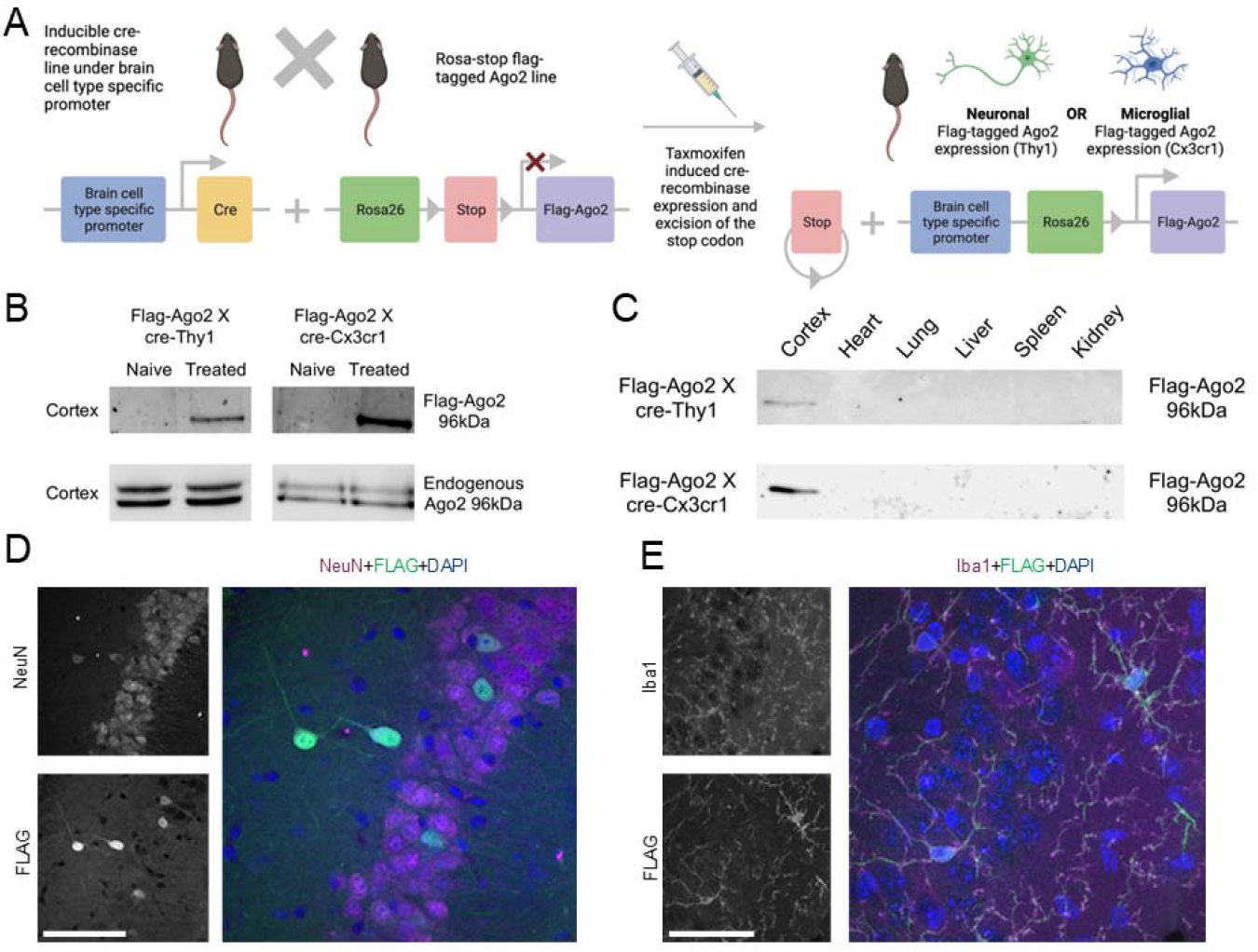
Cell type-specific FLAG-Ago2 mice. (A) Schematic showing the breeding scheme for generating mice expressing FLAG-Ago2 under a Thy1 or Cx3cr1 tamoxifen promoter. (B) Western blot analysis of Ago2 performed following FLAG immunoprecipitation from the two FLAG-Ago2 X cre lines confirming presence of the FLAG-tagged Ago2 protein following tamoxifen treatment. No FLAG-tagged Ago2 protein was detected in untreated controls. The wildtype Ago2 protein was present in the whole cortical lysate from the same animals. Representative images from n = 3 experiments. (C) Representative western blot using anti-Ago2 antibodies following a FLAG immunoprecipitation from the two different mouse lines after tamoxifen treatment. A band at the expected molecular weight of the FLAG-Ago2 protein (∼96 kDa) was detected in brain samples but not other organs (tissue was pooled n = 3/lane and results are representative of two independent experiments). (D) Representative confocal images of FLAG tag in hippocampal tissue sections from tamoxifen-treated FLAG-Ago2Thy1 mice. Sections were counter-stained with antibodies against NeuN. Larger image shows overlay of DAPI (nuclei). Note, scattered presence of FLAG in neurons. Scale bar, 100 µm. (E) Representative confocal images of FLAG tag in hippocampal tissue sections from tamoxifen-treated FLAG-Ago2Cx3cr1 mice. Sections were counter-stained with antibodies against microglia marker Iba1. Larger image shows overlay of DAPI (nuclei). Note, widespread appearance of FLAG in microglia. Scale bar, 66 µm.

Using western blotting, we first confirmed that tamoxifen treatment (+) of the resulting FLAG-Ago2Thy1 and FLAG-Ago2Cx3cr1 lines resulted in detectable FLAG at the expected molecular weight within the mouse brain (Figure 2B). In contrast, only the endogenous Ago2 was detected in cortical lysates from non-tamoxifen treated FLAG-Ago2Thy1 and FLAG-Ago2Cx3cr1 mice (Figure 2B). Next, we confirmed that the FLAG-tagged Ago2 from both lines was expressed in lysates from the brain of tamoxifen-treated mice but not a selection of other organs tested (Figure 2C).

We then used confocal microscopy to confirm that the FLAG-Ago2 was correctly restricted to either neurons (FLAG-Ago2Thy1+) or microglia (FLAG-Ago2Cx3cr1+). Hippocampal tissue sections from FLAG-Ago2Thy1+ mice were immunostained with antibodies against FLAG and neuronal nuclear protein (NeuN). Analysis of double-stained sections revealed expression of FLAG in a subset of neurons of various morphologies in the hippocampus (Figure 2D). This suggests that the FLAG-tagged Ago2 is likely expressed by both excitatory and inhibitory neurons, consistent with the expected labelling pattern using a *Thy1* promoter (Young et al., 2008).

Next, we stained sections from FLAG-Ago2Cx3cr1+ mice with antibodies against FLAG and the microglial marker Iba1. Staining for Iba1 had the expected appearance of microglia, with a small cell body from which multiple diffuse processes extended, and there was extensive co-localisation of FLAG staining with the microglial marker (Figure 2E). These experiments confirm the two different mice lines express neuronal and microglial cell type-restricted expression of FLAG-Ago2.

### Status epilepticus in cell type-specific FLAG-Ago2 mice

Macroscopic inspection of the brains of the FLAG-Ago2Thy1+ and FLAG-Ago2Cx3cr1+ mice revealed normal morphology (data not shown). Nevertheless, we next assessed the severity and effects of SE in the transgenic mice. Seizures were induced by intra-amygdala kainic acid and recorded using skull-mounted EEG, comparing the two different tamoxifen-treated FLAG-Ago2 lines to untreated transgenic lines.

Intra-amygdala kainic acid triggered prolonged seizures in all mice, characterised by high amplitude, high frequency discharges (Figure 3A). Semi-quantitative analysis of EEGs revealed there were no significant differences in seizure severity between the different mouse lines (Figure 3B). To extend these findings, we analysed SE-induced neuronal death in the ipsilateral hippocampus of a subset of the mice 24 h after SE. In wildtype mice, seizure-induced damage was observed within the ipsilateral CA3 subfield as expected (Mouri et al., 2008), whereas the bordering seizure-resistant CA2 subfield was spared (Figure 3C). Seizure-induced damage was similar for both FLAG-Ago2Thy1+ and FLAG-Ago2Cx3cr1+ mice and their untreated littermates subject to SE (Figure 3C). Together these data indicate that expression of the FLAG-Ago2 does not significantly alter the electrographic and pathologic features of the intra-amygdala kainic acid model.

**Figure 3.**
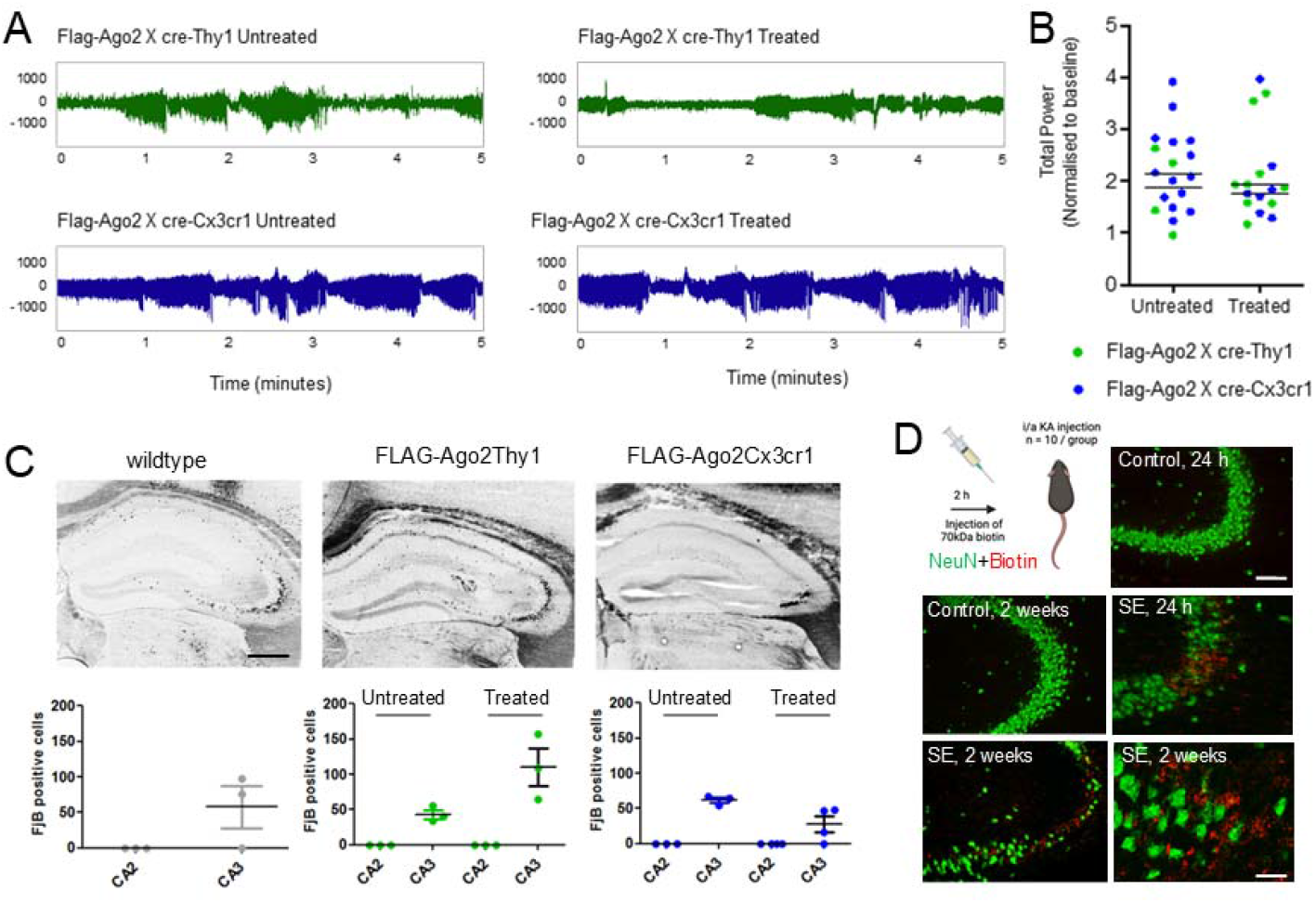
Effects of SE in FLAG-Ago2Thy1 and FLAG-Ago2Cx3cr1 mice. (A) Representative EEG recordings of SE following intraamygdala kainic acid in tamoxifen treated and untreated FLAG-Ago2Thy1 and FLAG-Ago2Cx3cr1 mice. (B) Analysis of EEG total power during SE in tamoxifen treated and untreated FLAG-Ago2Thy1 and FLAG-Ago2Cx3cr1 mice. No differences in seizure severity were observed between groups. (C) Representative images and counts of FluoroJade B positive cells (irreversible neuronal damage) observed in the damage-sensitive CA3 and damage-resistant CA2 subfields of the hippocampus. Similar seizure damage was observed in all groups. Scale bar, 500 µm. (D) Assessment of BBB permeability observed following SE in wildtype mice. Biotin-dextran amine (BDA) was injected 2 h or 11 days following intraamgydala injection. Injected mice were perfused 72 h later and staining performed to assess BBB disruption. Note, BDA co-localised with sites of neuronal damage in the CA3 hippocampal region. Scale bar, upper panel = 100 µm; lower panel 50 µm.

We next sought evidence that SE causes blood-brain barrier disruption in this model sufficient to allow passage of a large protein such as Ago2 to pass between blood and brain. For this, we injected wildtype mice with a biotin tracer, 2 hours after injection of kainic acid, a time-point when the blood-brain barrier had previously been found to be maximally disrupted in this model (Reschke et al., 2021) (Figure 3D). Staining tissue sections from mice injected with the tracer revealed staining within the CA3 subfield, the major site of damage in the model, but not in control mice (Figure 3D). Similar results were obtained from two-week post-SE mice. This indicates that changes to the integrity of the blood brain barrier in epilepsy are likely sufficient to allow direct passage of a protein the size of Ago2 into the blood.

### Brain cell type-specific origins of circulating Ago2-bound miRNAs

Last, we sought proof-of-principle that miRNAs from the brain can be detected associated with the FLAG-Ago2 in the plasma of mice after seizures. Here, we focused on miRNAs that were likely brain-enriched and potential biomarkers of TLE, selecting let-7i-5p (present data and (Lagos-Quintana et al., 2002)) and miR-19b-3p, which we previously identified as an Ago2-enriched, cerebrospinal fluid biomarker of TLE in humans (Raoof et al., 2017). We subjected FLAG-Ago2Thy1+ and FLAG-Ago2Cx3cr1+ mice to prolonged seizures induced by kainic acid and then immunoprecipitated FLAG from plasma collected 2 weeks later (Figure 4A). RNA was extracted from the eluted FLAG and subjected to miRNA assays for either let-7i-5p or miR-19b-3p (Figure 4B-E). We observed significantly elevated levels of Flag-Ago2-bound miR-19b-3p in both the FLAG-Ago2Thy1+ and FLAG-Ago2Cx3cr1+ mice which experienced SE compared to those which did not. Let7i-5p was elevated in FLAG-eluates from the FLAG-Ago2Cx3cr1+ which experienced SE but not the FLAG-Ago2Thy1+ mice (Figure 4B-E). These findings indicate that a portion of the circulating pool of miRNA has originated from the brain, including a contribution from both microglia and neurons.

**Figure 4.**
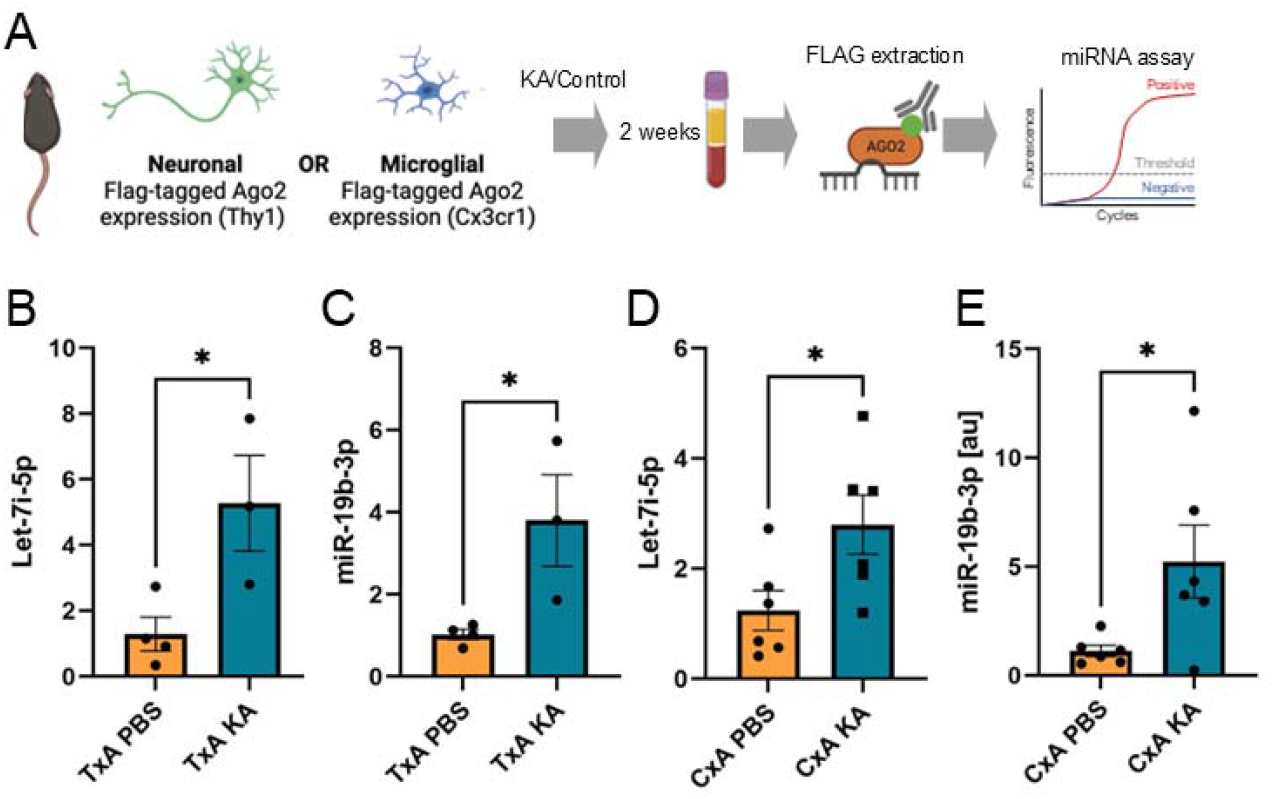
Detection of brain-originating FLAG-Ago2-carried miRNAs in plasma following SE in mice. (A) Schematic showing experimental design. Tamoxifen-treated FLAG-Ago2Thy1 and FLAG-Ago2Cx3cr1 mice were subject to SE (or control) and plasma sampled two weeks later. FLAG was immunoprecipitated from the plasma, RNA extracted and miRNAs measured. (B – E) Graphs show elevated levels of let-7i-5p and miR-19b-3p in FLAG extracts from the plasma of individual mice subject to SE compared to control. *P < 0.05.

## Discussion

The development of blood-based miRNA biomarkers for epilepsy would benefit from more direct evidence of pathophysiologic relevance. Here we report that known brain cell type-enriched miRNAs can be detected bound to Ago2 in the plasma of mice subjected to SE. By generating transgenic mice with a FLAG-tagged Ago2 restricted to one of two major subclasses of brain cells and then eluting miRNA from FLAG immunoprecipitates from plasma, we demonstrate that a portion of the circulating miRNA pool likely originates from the brain after SE. Taken together, these findings increase support for circulating miRNAs having mechanistic relevance as biomarkers of epilepsy.

There remains an unmet need for biomarkers to support diagnosis of epilepsy and recent or impending seizures. To this end, multiple studies have reported elevated levels of brain-enriched or inflammation-associated miRNAs in biofluids such as plasma in experimental and human epilepsy (Enright et al., 2018). The present study was designed to increase confidence that these circulating biomarkers have direct mechanistic links to the pathophysiology (Brindley et al., 2019). For example, the altered miRNA profiles in blood and other biofluids reported in patients might reflect non-CNS effects of the disease such as elevated inflammation or responses to anti-seizure medicines. The present study addressed this concern by providing evidence that miRNAs that were Ago2-loaded in the brain can be detected among the plasma-based miRNA biomarker signal.

Previous studies have confirmed that when expressed in the mouse brain, FLAG-Ago2 carries an extensive range of miRNAs representative of the functional miRNA pool (Tan et al., 2013;Brennan et al., 2020). Here, we adapted this approach to develop an *in vivo* model to test for brain cell-derived miRNAs in the circulation. Combining *in vivo* cre/loxP technology, immunoprecipitation methods and RT-qPCR, we placed FLAG-Ago2 expression under *Thy1* or *Cx3cr1* control and could elute miRNAs in FLAG pull-downs from the plasma of mice. Plasma levels of the tested FLAG-bound miRNAs were broadly elevated in both lines, consistent with increased release following seizures. Among those was let-7i, which is brain-enriched, although expressed by other organs (Keller et al., 2022). Hippocampal levels of let-7i decline after pilocarpine-induced SE (Risbud and Porter, 2013) but increase in the plasma after traumatic brain injury in rats (Balakathiresan et al., 2012), suggesting circulating let-7i may be a broad biomarker for epileptogenic brain injuries. Let-7i may also be a therapeutic target via effects on brain-derived neurotrophic factor (BDNF)-progesterone signalling and neuroprotective responses (Nguyen et al., 2018;He et al., 2022). A second miRNA, miR-19b showed a similar profile of elevation in FLAG-Ago2 from neuronal and microglial lines. Elevated levels of miR-19b have been reported in the cerebrospinal fluid of TLE and SE patients (Raoof et al., 2017) with functional studies indicating miR-19b controls glial signalling (Kalozoumi et al., 2018). Together, this supports there being a brain- and brain-cell type miRNA signature present in the circulation, increasing confidence that circulating miRNA have direct pathophysiologic links to events in the brain in experimental epilepsy. Further optimisation to elute microRNAs from low-abundance FLAG-Ago2 in plasma could enable a complete profile of the circulating miRNAs derived from these brain cell types.

The ipsilateral CA3 subfield displays neurodegeneration, chronic BBB disruption and generates the spontaneous seizures in the present model (Li et al., 2008;Reschke et al., 2021). Accordingly, the hippocampus may be the source of the miRNAs eluted along with FLAG-Ago2 from plasma. Indeed, the tamoxifen-inducible *Thy1* promoter restricted the expression of FLAG-Ago2 to neurons, including hippocampal CA subfield neurons. We observed mixed neuronal populations express FLAG-Ago2, as expected (Young et al., 2008). Thus, we cannot assign plasma-detected miRNAs to specific affected brain structures or neuronal subtypes. The sparse expression of FLAG-Ago2 in neurons suggests we are likely to have underestimated the full contingent of neuron-derived miRNAs in the plasma. The detection of FLAG-Ago2-bound miR-19b-3p in the *Cx3cr1*-driven line indicates a portion of the circulating miRNA pool may come from non-neuronal cells, likely microglia. This supports the multiple brain cell type contribution implied by our Ago2 sequencing of plasma from mice after SE. The finding is also consistent with active miRNA signalling by microglia in epilepsy (Brennan and Henshall, 2020). Indeed, microglia are increasingly understood to perform complex functions in the brain, enhancing inhibitory neurotransmission (Badimon et al., 2020), but also promoting a pro-epileptogenic inflammatory environment by engaging incoming immune cells (Kumar et al., 2022). The specificity of *Cx3cr1* for microglia is incomplete, and we cannot exclude a non-brain-resident macrophage contribution to the miRNA eluted from the FLAG-Ago2 mice using this promoter (Yona et al., 2013). Nevertheless, immune cell entry occurs in the epileptogenic focus in drug-resistant epilepsy (Kumar et al., 2022), and the peripherally-detected FLAG-Ago2 from the *Cx3cr1* mouse line containing miRNA may have originated from centrally-driven pathophysiologic actions of these cells.

Additional mouse lines could be generated to further explore brain region or cell type-specific contributions to the circulating biomarker signal. Astrocytes represent a key cell type implicated in epileptogenesis, including in this model (Li et al., 2008), and where miRNA-driven signalling is established (Simonato et al., 2021). Our attempted use of a *Gfap* promoter to drive a FLAG-Ago2 astrocyte line was unsuccessful. Single cell and other data on cell specification markers could yield alternative approaches (Andersson et al., 2014). Other cell types, including oligodendroglia, are increasingly linked to the pathogenesis of epilepsy and progressive white matter injury and miRNA from these sources may contribute to the circulating miRNA pool (Dubey et al., 2022).

Ago2 sequencing provided broad support for a significant contribution of brain-originated miRNAs to the circulating miRNAs pool. We found higher levels of multiple brain-enriched miRNAs in the plasma of mice within 4 h of SE, which declined at later time points. This likely reflects the early and transient large-scale opening of the BBB in the hours following SE in the model (Michalak et al., 2013;Reschke et al., 2021). Brain-derived miRNAs continued to be elevated, however, in samples up to two weeks after SE. This could be due to ongoing spontaneous seizure-induced breaches in BBB integrity (Reschke et al., 2021;Greene et al., 2022). We do not know the mechanism of passage of brain miRNAs into the circulation. The abundance of the presumed brain-originated miRNAs in the plasma largely match the relative abundances of neurons and glia in the brain. It is unknown whether cell lysis or pathway-directed release of Ago2-bound miRNAs contributes most to the signal in the circulation. This could be resolved using additional and complementary methods to label specific brain cell proteins or microvesicles (e.g. exosomes) prior to subsequent detection in the circulation (Parker and Pratt, 2020).

AGO2/Ago2 is considered the main catalytic engine of the RISC but other Ago proteins are involved in miRNA-based gene silencing (Czech and Hannon, 2011). The tissue distribution and function of different Ago proteins varies and this could influence the biomarker signal we obtained in plasma (Turchinovich and Burwinkel, 2012). Here we focused on Ago2-carried miRNA in the circulation. Future studies could profile the circulating miRNA content of other Ago isoforms to determine if one or more contain a signal with highest sensitivity and specificity to distinguish epilepsy samples from controls. A limitation of the present study was the lack of biological replicates for the sequencing part of the study, due to the necessarily large number of mouse samples that had to be pooled to create a sample that could yield enough Ago2-bound small RNA to be sequenced. Advances in using lower input RNA may enable replicates to be generated from individual animals (Dave et al., 2019). This would also allow plasma miRNA profiles to be aligned to individual electrophysiological or neuropathological profiles. Comparing the Ago pool to the total pool or the miRNA enclosed in microvesicles may yield further improvements in signal-to-noise that could guide decisions on technology for point-of-care testing (Enright et al., 2018;Brindley et al., 2019). Thus, the present findings may have implications for how we process plasma for biomarker studies (Dave et al., 2019). If the Ago2-bound signal carries the superior diagnostic yield, then measurement assays may need to be adapted to include a step to select for this protein-conveyed miRNA population. Notably, there are rapid, non-amplified miRNA detection technologies that have promise as point-of-care devices but these currently use whole plasma and so would need an adaptation to recover the Ago fraction (McArdle et al., 2017;Dave et al., 2019).

Clinical studies have identified circulating miRNA biomarkers of epilepsy in CSF (Raoof et al., 2017) and blood (serum and plasma) (Wang et al., 2015;Enright et al., 2018;Raoof et al., 2018;Brennan et al., 2020;Ioriatti et al., 2020). Several of the same human epilepsy circulating miRNAs were detected here, bound to Ago2 in the plasma of mice. This supports the clinical relevance of the mouse model, which is refractory to various anti-seizure medicines (West et al., 2022), as well as the likely brain origins of some portion of the proposed miRNA biomarkers in clinical studies. However, we also identified miRNAs dysregulated in the Ago2 sequencing study which have not yet been reported in clinical studies. This suggests analysing specific pools of miRNA may uncover more subtle disease-associated molecular signatures and provide better diagnostic power than whole plasma analysis.

In summary, the present study demonstrates that a portion of the circulating miRNA signature in the blood is derived from the brain in experimental TLE. Brain cell type restricted expression of FLAG-Ago2 and subsequent FLAG immunoprecipitations indicate this includes neuronal and non-neuronal cell, namely microglial, contributions. These findings support ongoing efforts to develop miRNA-based biomarker panels to support diagnosis of epilepsy and seizures and may inform the design of technologies for their detection.

## Methods

### Animal studies

All animal experiments were performed in accordance with the European Union Directive (2010/63/EU) and were reviewed by the HPRA (19127/P001) and the Research Ethics Committee, RCSI (REC 842). Adult male C57BL/6JOlaHsd mice were bred in house in the Biomedical Research Facility (BRF) in RCSI unless otherwise indicated. All animals were housed 2-5/cage in the climate-controlled environment on a 12 h light/dark cycle. Food and water were provided *ab libitum*.

### Generation of cell-type specific FLAG-tagged-Argonaute-2 mice

Heterozygous transgenic mice expressing Rosa-Stop^fl/fl^-Flag-Ago2 were kindly provided by Anne Schaefer, Icahn School of Medicine, NYC (Schaefer et al., 2010;Tan et al., 2013). Homozygous FLAG-tagged Ago2 mice were crossed with inducible cre-recombinase mouse lines driven by brain cell type restricted promoters. Upon treatment with the inducing agent, tamoxifen, neuron (Thy1, Stock 012708) and microglia (Cx3cr1, Stock 021160) restricted expression of FLAG-tagged Ago2 resulted. New mouse lines produced as part of this study have genotypes *Thy1-cre^tg/+^;Rosa-Stop^fl/fl^-Flag-Ago2* and *Cx3cr1-cre^tg/tg^;Rosa-Stop^fl/fl^-Flag-Ago2.* Tamoxifen (0.1 mL of 10 mg/mL solution, Sigma-Aldrich, Ireland) was administered via intraperitoneal injection as follows; once a day for five days, two day rest, then once a day for five days.

### Murine model of SE, plasma collection and processing

SE was induced as previously described (Reschke et al., 2021) with some minor modifications. Briefly, adult C57/BL/6 mice were anesthetized. A guide cannula and three surface EEG electrodes were implanted. Kainic acid (0.3 μg) was injected into the ipsilateral amygdala in awake mice to trigger SE. To reduce mortality and morbidity, midazolam or lorazepam (8 mg/kg) was administered after 40 minutes (Diviney et al., 2015). Blood was collected from mice using the submandibular bleed technique (Brennan et al., 2018), into a 0.5 mL tube coated with K2 EDTA (0.5M; Sigma-Aldrich, Ireland) at various time points following SE. Samples were centrifuged at 1300 x g for 10 mins at 4°C to separate the plasma from blood cells. The plasma supernatant was removed and placed in a fresh tube and immediately placed on ice. Haemolysis levels were evaluated using a Nanodrop 2000 spectrophotometer and only samples with a reading of <0.25Abs at 414nm were used (Brennan et al., 2018).

### Extraction of small RNA from plasma fractions

Ago2 protein was extracted as described previously (Raoof et al., 2018). Briefly, pre-cleared plasma was incubated with Ago2 antibody (C34C6; Cell Signalling Technology) overnight at 4°C. Protein A/G beads were added, and samples were incubated at 4°C for 2 h. Bead-Ago2 complexes were pelleted by centrifugation, washed and RNA was extracted using the phenol-chloroform technique. 50 µL of plasma from 10 mice were pooled (final volume 500 µL) and the Ago2-bound fraction was extracted as outlined above. The FLAG-tagged Ago2 protein was precipitated from transgenic mouse plasma using the technique outlined above except the pre-cleared plasma was incubated with 10 µL of the Anti-FLAG M2 Magnetic Beads (Cat#: M8823 Sigma-Aldrich, Ireland) overnight at 4°C.

### Small RNA sequencing

The time-points for Ago2 sequencing were 4 h, 8 h, 24 h and 2 weeks after SE. Each SE sample was compared to a plasma pool collected from a control (intraamygdala PBS) mouse from the sample time point (n = 10/each). An exception was the 2 week time-point where the control sample failed to generate a library that passed QC and therefore the 2 week SE sample was compared to the 24 h control. Small RNA libraries from plasma were prepared using the Illumina TruSeq Small RNA Library Prep Kit (Illumina, UK) with reagent volumes halved to account for lower concentrations of miRNA to prevent excessive formation of adapter dimers. Small RNA libraries were separated by electrophoresis on Pippin Prep (Sage Science, US) using a 3% agarose gel cassette (Lab Tech, UK), selecting for RNA in the size range of 90-250 bp (adapted-ligated miRNA). Libraries were sequenced on an Illumina HiSeq.

### Bioinformatic analysis

FastQ files were downloaded from the Illumina BaseSpace website (basespace.illumina.com). The Chimera software was used to trim adapters, size-select miRNA reads and to align identified reads to the reference genome of interest. Reads are aligned to miRBase using BLASTn. This software allows 2-mismatched nucleotides. Downstream analyses to identify differentially expressed miRNA were performed using edgeR and Limma packages from Bioconductor. Graphics were produced using the glimma package. For mouse analysis, based on the fact that that the mice are genetically identical, a square root value (BCV) of 0.1 was set to allow an estimated p value based on dispersion values.

### Identification of brain and brain cell-type specific miRNAs

An extensive literature search was performed to identify reported enrichment of miRNA identified by RNAseq. The entire list of miRNAs that mapped to the mouse miRBase V21 database was compared against those reportedly enriched in different organs and brain regions (Lagos-Quintana et al., 2002) and enriched in different brain cell types *in vitro* (Jovicic et al., 2013) and *in vivo* (Butovsky et al., 2014). miRNAs were searched in PubMed to identify *in situ* hybridization studies (Schratt et al., 2006;Thompson et al., 2007;Cheng et al., 2009;Huse et al., 2009;Pena et al., 2009;Aronica et al., 2010;Nelson et al., 2010;Iyer et al., 2012;Ouyang et al., 2013;Kaalund et al., 2014;van Scheppingen et al., 2016) demonstrating colocalization of miRNAs with known brain cell type markers and using techniques such as miRAP (He et al., 2012).

### Small-scale individual Taqman assay

Brain cell type of origin was investigated using Taqman small-scale miRNA assay (Applied Biosystems) as described. Plasma from two male animals of the same genotype were pooled per sample and all amplifications were performed in triplicate in a 96-well plate and the comparative Ct values were measured on the 12K-Flex.

### Western blotting

Tissue samples were homogenised in 500 µL protein lysis buffer (0.1M NaCl, 20mM Tris-HCl, pH7.6, 1mM EDTA, pH8.0, 1% NP-40). Samples were centrifuged to pellet any cellular debris. FLAG-Ago2 was immunoprecipitated from 800 µg of tissue lysate using an Anti-FLAG M2 coated 96-well plates (P2983, Sigma-Aldrich, Ireland) as per the manufacturer’s guidelines. Lysates were removed and protein antibody-conjugates linked to the bottom of the plate were washed three times with 300 µL of ELISA wash buffer (100mM Tris buffered saline, pH8.2, 2mM MgCl_2_, 0.5% Tween-20 – all Sigma-Aldrich, Ireland). Protein was eluted by incubating wells in 2 X Laemelli buffer for 10 mins, shaking at 900rpm. FLAG-Ago2 immunoprecipitates and 30 µg of whole brain lysate were analysed using standard western blotting techniques using antibodies against FLAG and Ago2.

### Immunohistochemistry

Paraformaldehyde-fixed, free-floating sections were blocked in 10% normal goat serum, before incubation in antibodies against FLAG, NeuN and Iba1 overnight at 4°C. Sections were washed and in suitable secondary antibody (Alexa Fluor 488 or Alexa Fluor 568) diluted in PBS containing 0.1% Triton-X 100 for 2 h at room temperature. Sections were investigated under Leica LSM confocal microscope at 40X magnification.

### FluoroJade B staining

Fluorojade B staining was performed to assess SE-induced neuronal death (Mouri et al., 2008). Briefly, brain slices were cryosectioned at a thickness of 12 μm and tissue was fixed in 4% paraformaldehyde. Slices were rehydrated in ethanol, incubated in a 0.006% potassium permanganate and finally incubated in 0.001% Fluorojade B. Neuronal damaged was assessed by manually counting of Flurojade B positive cells in the CA1, CA2, CA3 and dentate gyrus hippocampal regions under a Leica Nikon 2000s epifluorescence microscope.

### Biotin administration and staining

BBB disruption was assessed by systemically injecting Biotin dextran amine (BDA, 70 kDa) following SE and staining for the presence of BDA in the mouse brain. For the acute experiments, BDA at a concentration of 90 mg/kg was administered via intraperitonial injection 2 h following KA injection. Then, 72 h following BDA injection, mice were PBS perfused and brains were snap frozen. To confirm disruption to the BBB in the chronic phase, BDA was injected 11 days after SE and then brains collected 72 h later. Mouse brain slices were cryosectioned at a thickness of 12 μm and mounted on slides and formalin fixed. Staining was performed as above, however streptavidin-AF594 (1:500) was added to the secondary antibody and incubated 2 h at room temperature. Sections were investigated under a Leica Nikon 2000s epifluorescence microscope.

## Acknowledgments

We thank Anne Schaefer for the generous donation of Rosa-Stop^fl/fl^-Flag-Ago2 mice. We thank the operations team at FutureNeuro for administrative support and the staff at the biomedical research facilities at RCSI. This publication has emanated from research supported by a research grant from Science Foundation Ireland (SFI) under agreement 16/RC/3948 (FutureNeuro) and 13/IA/1891, the European Union’s “Seventh Framework” Programme (FP7) under Grant Agreement 602130 (EpimiRNA), Horizon Europe 2020 awards EpimiRGen and EpimiRTherapy, the Wellcome Trust (222648/Z/21/Z), and the Higher Education Authority (SeeDeepER).

