## Supplementary File for "Brain cell-specific origin of circulating microRNA biomarkers in experimental temporal lobe epilepsy"

**Supplementary Figure S1**

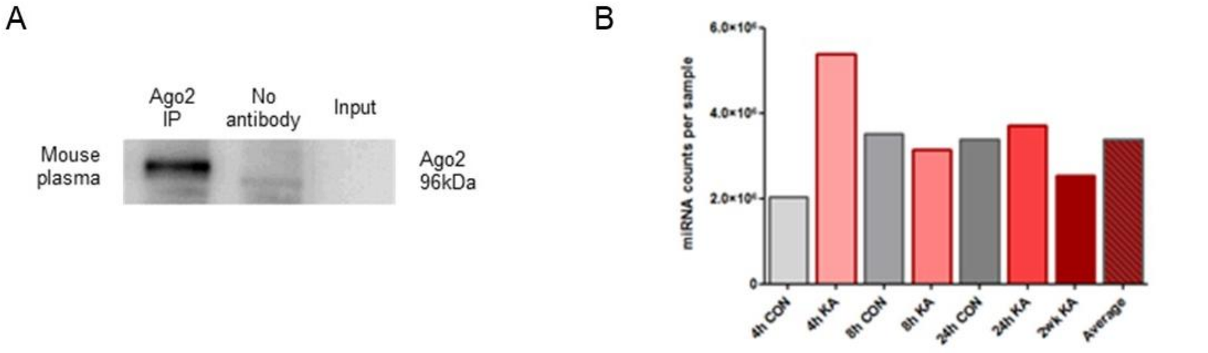


**Supplementary Figure S1** *Ago elution from plasma and small RNAseq reads across samples* (A) Immunoblot confirming elution of Ago2 from plasma. Alongside the immunoprecipitations, a control containing no antibody and the supernatant (remainder of the sample, once the Ago2 protein was precipitated from the sample were also analysed). (B) Total normalised counts per million present in at time points (4 h, 8 h, 24 h and 2 wk-post PBS/KA injection).

**Supplementary Figure S2**

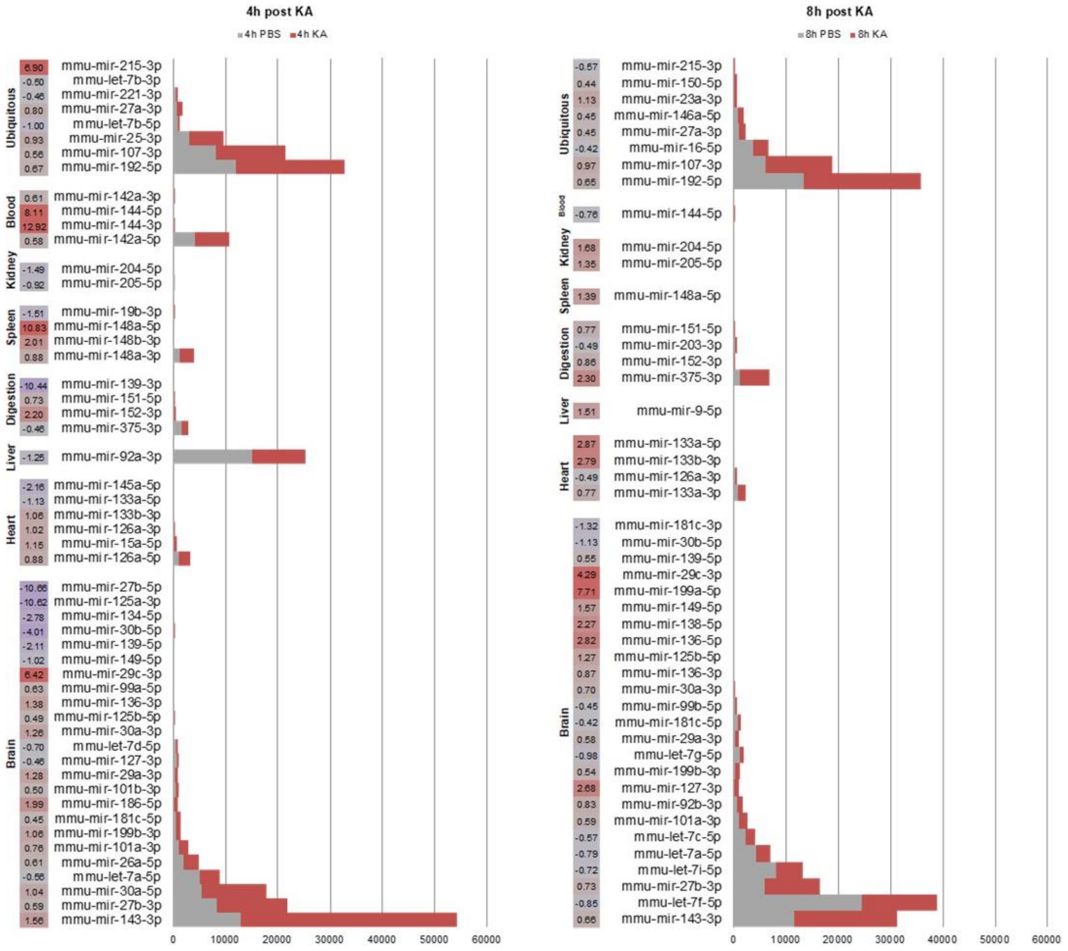


**Supplementary Figure S2.** *Circulating miRNA changes following SE in mice.* Normalised counts of miRNA detected at 4 h and 8 h time points reportedly enriched in different tissue types; brain, heart, liver, digestive system, kidney, blood and ubiquitously expressed miRNAs were graphed. Log2 fold change differences included alongside counts and enrichment. MiRNAs with 50 counts per million (CPM) or more in PBS or KA samples at a time point were considered present for the organ enrichment portion of this analysis. MiRNAs with a p value of <0.05 and a fold change of +/- 1.5 were considered differentially regulated.

**Supplementary Figure S3**

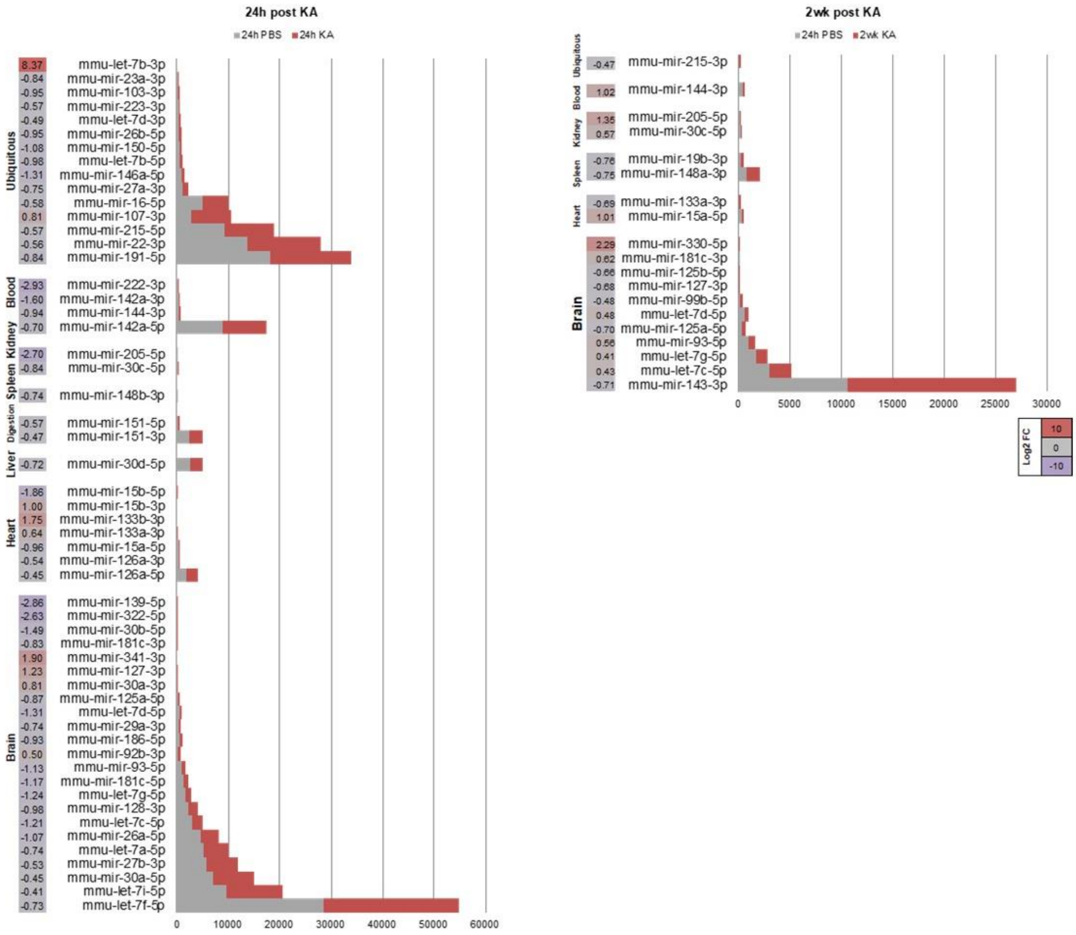


**Supplementary Figure S3** *Circulating miRNA changes following SE in mice.* Normalised counts of miNA detected at 24 h and 2 week time points reportedly enriched in different tissue types; brain, heart, liver, digestive system, kidney, blood and ubiquitously expressed miRNAs were graphed. Log2 fold change differences included alongside counts and enrichment. MiRNAs with 50 counts per million (CPM) or more in PBS or KA samples at a time point were considered present for the organ enrichment portion of this analysis. MiRNAs with a p value of <0.05 and a fold change of +/-1.5 were considered differentially regulated.

**Supplementary Table S1** Sources of information on miRNA tissue and organ enrichment

| miRNA | Tissue | Reference | Brain cell | Reference |
|---|---|---|---|---|
| **Let-7a-5p** | Brain | (Ludwig et al., 2016) | Neuron | (Wang et al., 2014) |
| **Let-7b-3p** | Ubiquitous | (Bargaje et al., 2010) | | |
| **Let-7b-5p** | Ubiquitous | (Bargaje et al., 2010) | Neuron | (Butovsky et al., 2014) |
| **Let-7c-5p** | Brain | (Ludwig et al., 2016) | Neuron | (Butovsky et al., 2014) |
| **Let-7d-3p** | Ubiquitous | (Ludwig et al., 2016, Lagos-Quintana et al., 2001) | | |
| **Let-7d-5p** | Brain | (Ludwig et al., 2016, Lagos-Quintana et al., 2001) | Neuron | (Butovsky et al., 2014) |
| **Let-7f-5p** | Brain | (Ludwig et al., 2016, Lagos-Quintana et al., 2001) | | |
| **Let-7g-5p** | Brain | (Lagos-Quintana et al., 2001) | Microglia | (Butovsky et al., 2014) |
| **Let-7i-5p** | Brain | Lagos-Quintana et al., 2001) | Neuron | (Butovsky et al., 2014) |
| **miR-100-5p** | | | Astrocyte | (Butovsky et al., 2014) |
| **miR-101a-3p** | Brain | Lagos-Quintana et al., 2001) | | |
| **miR-101b-3p** | Brain | Lagos-Quintana et al., 2001) | | |
| **miR-101c** | Brain | Lagos-Quintana et al., 2001) | | |
| **miR-103-3p** | Ubiquitous | (Bargaje et al., 2010) | Brain | (Wang et al., 2014 |
| **miR-106a-5p** | | | Microglia | (Butovsky et al., 2014) |
| **miR-107-3p** | Ubiquitous | (Bargaje et al., 2010) | Neuron | (Nelson et al., 2006) |
| **miR-10a-5p** | Ubiquitous | (Ludwig et al., 2016, Guo et al., 2014, Liang et al., 2007) | | |
| **miR-1191a** | | | Neuron | (Butovsky et al., 2014) |
| **miR-124-3p** | | | Neuron | (Butovsky et al., 2014) |
| **miR-124-5p** | | | Neuron | (Jovicic et al., 2013) |
| **miR-125a-3p** | Brain | (Guo et al., 2014, Lagos-Quintana et al., 2001) | | |
| **miR-125a-5p** | Brain | (Guo et al., 2014, Lagos-Quintana et al., 2001) | Neuron | (Butovsky et al., 2014) |
| **miR-125b-5p** | Brain | (Ludwig et al., 2016) | | |
| **miR-126a-3p** | Heart | (Guo et al., 2014, Landgraf et al., 2007) | Astrocyte | (Butovsky et al., 2014) |
| **miR-126a-5p** | Heart | (Guo et al., 2014, Landgraf et al., 2007) | Astrocyte | (Butovsky et al., 2014) |
| **miR-127-3p** | Brain | (Ludwig et al., 2016) | Neuron | (Butovsky et al., 2014) |
| **miR-128-3p** | Brain | (Lagos-Quintana et al., 2002) | Neuron | (Tan et al., 2013) |
| **miR-129-2-3p** | | | Neuron | (Jovicic et al., 2013) |
| **miR-129-5p** | | | Neuron | (Jovicic et al., 2013) |
| **miR-130a-3p** | | | Astrocyte | (Butovsky et al., 2014) |
| **miR-132-3p** | Brain | (Lagos-Quintana et al., 2002) | Neuron | (Hansen et al., 2013) (Thompson et al., 2007) |
| **miR-133a-3p** | Heart | (Bargaje et al., 2010) | Oligodendrocyte | (Butovsky et al., 2014) |
| **miR-133a-5p** | Heart | (Ludwig et al., 2016, Guo et al., 2014, Lee et al., 2008, Bargaje et al., 2010) | | |
| **miR-133b-3p** | Heart | (Ludwig et al., 2016, Guo et al., 2014, Lee et al., 2008, Bargaje et al., 2010) | Neuron | (He et al., 2012) |
| **miR-134-5p** | Brain | (Lagos-Quintana et al., 2002) | Neuron | (Jimenez-Mateos et al., 2012) |
| **miR-136-3p** | Brain | (Lagos-Quintana et al., 2002) | Neuron | (Jovicic et al., 2013) |
| **miR-136-5p** | Brain | (Lagos-Quintana et al., 2002) | Neuron | (Butovsky et al., 2014) |
| **miR-137-3p** | | | Neuron | (Butovsky et al., 2014) |
| **miR-138-5p** | Brain | (Guo et al., 2014) | Neuron | (Krol et al., 2010) |

| | | | | |
|---|---|---|---|---|
| **miR-139-3p** | Digestion | (Ludwig et al., 2016) | | |
| **miR-139-5p** | Brain | (Ludwig et al., 2016) | Neuron | (Jovicic et al., 2013) |
| **miR-140-3p** | Digestion | Lagos-Quintana et al., 2002) | | |
| **miR-142a-3p** | Blood | (Landgraf et al., 2007, Ludwig et al., 2016) | Microglia | (Butovsky et al., 2014) |
| **miR-142a-5p** | Blood | (Landgraf et al., 2007, Ludwig et al., 2016) | Microglia | (Butovsky et al., 2014) |
| **miR-143-3p** | Brain | (Liang et al., 2007) | Astrocyte | (Jovicic et al., 2013) |
| **miR-144-3p** | Blood | (Ludwig et al., 2016) | | |
| **miR-144-5p** | Blood | (Ludwig et al., 2016) | | |
| **miR-145a-5p** | Heart | (Lagos-Quintana et al., 2001) | Astrocyte | (Butovsky et al., 2014) |
| **miR-146a-5p** | Ubiquitous | (Guo et al., 2014, Landgraf et al., 2007, Bargaje et al., 2010) | Astrocyte | (Iyer et al., 2012) |
| **miR-148a-3p** | Spleen | (Lagos-Quintana et al., 2001) | Neuron | (Butovsky et al., 2014) |
| **miR-148a-5p** | Spleen | (Lagos-Quintana et al., 2001) | | |
| **miR-148b-3p** | Spleen | (Lagos-Quintana et al., 2001) | | |
| **miR-149-5p** | Brain | (Ludwig et al., 2016, Guo et al., 2014) | | |
| **miR-150-5p** | Ubiquitous | (Ludwig et al., 2016, Landgraf et al., 2007) | Microglia | (Jovicic et al., 2013) |
| **miR-151-3p** | Digestion | (Lagos-Quintana et al., 2001) | | |
| **miR-151-5p** | Digestion | (Lagos-Quintana et al., 2001) | Astrocyte | (Butovsky et al., 2014) |
| **miR-152-3p** | Digestion | (Lagos-Quintana et al., 2001) | | |
| **miR-154-5p** | | | Neuron | (Jovicic et al., 2013) |
| **miR-15a-5p** | Heart | (Lagos-Quintana et al., 2001) | Microglia | (Butovsky et al., 2014) |
| **miR-15b-3p** | Heart | (Lagos-Quintana et al., 2001) | | |
| **miR-15b-5p** | Heart | (Lagos-Quintana et al., 2001) | Neuron | (Natera-Naranjo et al., 2010) |
| **miR-16-5p** | Ubiquitous | (Lagos-Quintana et al., 2001) | Microglia | (Butovsky et al., 2014) |
| **miR-181a-5p** | | | Microglia | (Butovsky et al., 2014) |
| **miR-181b-5p** | | | Oligodendrocyte | Lau (Lau et al., 2008) |
| **miR-181c-3p** | Brain | (Lee et al., 2008) | | |
| **miR-181c-5p** | Brain | (Lee et al., 2008) | | |
| **miR-182-5p** | | | Neuron | (Smrt et al., 2010) |
| **miR-184-3p** | | | Astrocyte | (Butovsky et al., 2014) |
| **miR-185-5p** | | | Neuron | (Smrt et al., 2010) |
| **miR-186-5p** | Brain | (Guo et al., 2014) | | |
| **miR-188-5p** | | | Neuron | (Jovicic et al., 2013) |
| **miR-191-5p** | Ubiquitous | (Bargaje et al., 2010) | Microglia | (Butovsky et al., 2014) |
| **miR-192-5p** | Ubiquitous | (Guo et al., 2014, Landgraf et al., 2007, Liang et al., 2007) | | |
| **miR-193a-3p** | | | Astrocyte | (Jovicic et al., 2013) |
| **miR-1983** | | | Neuron | (Butovsky et al., 2014) |
| **miR-199a-5p** | Brain | (Guo et al., 2014, Liang et al., 2007) | | |
| **miR-199b-3p** | Brain | (Guo et al., 2014, Liang et al., 2007) | | |
| **miR-19a-3p** | | | Microglia | (Butovsky et al., 2014) |
| **miR-19b-3p** | Spleen | (Lagos-Quintana et al., 2001) | Neuron | (Natera-Naranjo et al., 2010) |
| **miR-203-3p** | Digestion | (Liang et al., 2007, Lee et al., 2008, Bargaje et al., 2010) | | |
| **miR-204-5p** | Kidney | (Guo et al., 2014, Lee et al., 2008, Bargaje et al., 2010) | Neuron | (Butovsky et al., 2014) |
| **miR-205-5p** | Kidney | (Ludwig et al., 2016) | | |

| | | | | |
|---|---|---|---|---|
| **miR-20a-5p** | | | Oligodendrocyte | (Butovsky et al., 2014) |
| **miR-210-3p** | | | Astrocyte | (Jovicic et al., 2013) |
| **miR-215-3p** | Ubiquitous | (Lee et al., 2008, Bargaje et al., 2010) | | |
| **miR-215-5p** | Ubiquitous | (Lee et al., 2008, Bargaje et al., 2010) | | |
| **miR-216b-5p** | | | Oligodendrocyte | (Butovsky et al., 2014) |
| **miR-218-5p** | | | Neuron | (Butovsky et al., 2014) |
| **miR-21a-5p** | | | Astrocyte | (Bhalala et al., 2012) |
| **miR-221-3p** | Ubiquitous | Bargaje (Bargaje et al., 2010) | Astrocyte | (Jovicic et al., 2013) |
| **miR-222-3p** | Blood | (Ludwig et al., 2016) | Astrocyte | (Jovicic et al., 2013) |
| **miR-223-3p** | Ubiquitous | (Guo et al., 2014, Landgraf et al., 2007, Lee et al., 2008, Bargaje et al., 2010) (Ludwig et al., 2016) | Astrocyte | (Jovicic et al., 2013) |
| **miR-22-3p** | Ubiquitous | (Lagos-Quintana et al., 2001) | Neuron | (Butovsky et al., 2014) |
| **miR-23a-3p** | Ubiquitous | (Lagos-Quintana et al., 2001, Bargaje et al., 2010) | Microglia | (Butovsky et al., 2014) |
| **miR-23b-3p** | | | Neuron | (Butovsky et al., 2014) |
| **miR-25-3p** | Ubiquitous | (Bargaje et al., 2010) | Microglia | (Butovsky et al., 2014) |
| **miR-26a-5p** | Brain | (Lagos-Quintana et al., 2001) | Neuron | (Pena et al., 2009) |
| **miR-26b-5p** | Ubiquitous | (Bargaje et al., 2010) | Neuron | (Butovsky et al., 2014) |
| **miR-27a-3p** | Ubiquitous | (Lagos-Quintana et al., 2001) | Microglia | (Butovsky et al., 2014) |
| **miR-27b-3p** | Brain | (Lagos-Quintana et al., 2001) | Oligodendrocyte | (Letzen et al., 2010) |
| **miR-27b-5p** | Brain | (Lagos-Quintana et al., 2001) | | |
| **miR-28a-5p** | | | Oligodendrocyte | (Butovsky et al., 2014) |
| **miR-29a-3p** | Brain | (Lagos-Quintana et al., 2001) | Astrocyte | (Butovsky et al., 2014) |
| **miR-29b-3p** | | | Microglia | (Butovsky et al., 2014) |
| **miR-29c-3p** | Brain | (Lagos-Quintana et al., 2001) | Astrocyte | (Butovsky et al., 2014) |
| **miR-300-3p** | | | Neuron | (Jovicic et al., 2013) |
| **miR-3099-3p** | | | Astrocyte | (Butovsky et al., 2014) |
| **miR-30a-3p** | Brain | (Lagos-Quintana et al., 2001) | | |
| **miR-30a-5p** | Brain | (Lagos-Quintana et al., 2001) | | |
| **miR-30b-5p** | Brain | (Lagos-Quintana et al., 2001) | Neuron | (Butovsky et al., 2014) |
| **miR-30c-5p** | Kidney | (Bargaje et al., 2010) | Astrocyte | (Butovsky et al., 2014) |
| **miR-30d-5p** | Liver | (Lagos-Quintana et al., 2001) | Astrocyte | (Butovsky et al., 2014) |
| **miR-31-5p** | | | Astrocyte | (Jovicic et al., 2013) |
| **miR-320-3p** | | | Neuron | (Wang et al., 2014 |
| **miR-322-3p** | | | Oligodendrocyte | (Jovicic et al., 2013) |
| **miR-322-5p** | Brain | (Ludwig et al., 2016) | | |
| **miR-330-5p** | Brain | (Liang et al., 2007, Guo et al., 2014) | | |
| **miR-335-5p** | | | Neuron | (Jovicic et al., 2013) |
| **miR-337-3p** | | | Neuron | (Jovicic et al., 2013) |
| **miR-338-3p** | | | Oligodendrocyte | (Butovsky et al., 2014) |
| **miR-338-5p** | | | Neuron | (Jovicic et al., 2013) |
| **miR-340-3p** | | | Astrocyte | (Butovsky et al., 2014) |
| **miR-340-5p** | Brain | (Lee et al., 2008) | Microglia | (Butovsky et al., 2014) |
| **miR-341-3p** | Brain | (Lee et al., 2008) | Neuron | (Jovicic et al., 2013) |
| **miR-34c-5p** | | | Neuron | (Butovsky et al., 2014) |

| | | | | |
|---|---|---|---|---|
| **miR-350-3p** | | | Microglia | (Butovsky et al., 2014) |
| **miR-351-5p** | | | Oligodendrocyte | (Jovicic et al., 2013) |
| **miR-369-3p** | | | Neuron | (Butovsky et al., 2014) |
| **miR-369-5p** | | | Neuron | (Jovicic et al., 2013) |
| **miR-375-3p** | Digestion | (Liang et al., 2007) | | |
| **miR-376b-3p** | | | Neuron | (Jovicic et al., 2013) |
| **miR-410-3p** | | | Neuron | (Jovicic et al., 2013) |
| **miR-411-5p** | | | Neuron | (Jovicic et al., 2013) |
| **miR-423-3p** | | | Oligodendrocyte | (Butovsky et al., 2014) |
| **miR-431-5p** | | | Neuron | (Jovicic et al., 2013) |
| **miR-433-3p** | | | Neuron | (Butovsky et al., 2014) |
| **miR-434-3p** | | | Neuron | (Butovsky et al., 2014) |
| **miR-449a-5p** | | | Neuron | (Jovicic et al., 2013) |
| **miR-450a-5p** | | | Oligodendrocyte | (Jovicic et al., 2013) |
| **miR-451a** | | | Astrocyte | (Raoof et al., 2017) |
| **miR-485-5p** | | | Neuron | (Jovicic et al., 2013) |
| **miR-497a-5p** | | | Oligodendrocyte | (Butovsky et al., 2014) |
| **miR-503-5p** | | | Oligodendrocyte | (Jovicic et al., 2013) |
| **miR-532-5p** | | | Oligodendrocyte | (Butovsky et al., 2014) |
| **miR-541-5p** | | | Neuron | (Jovicic et al., 2013) |
| **miR-542-3p** | | | Oligodendrocyte | (Jovicic et al., 2013) |
| **miR-653-5p** | | | Oligodendrocyte | (Butovsky et al., 2014) |
| **miR-669a-5p** | | | Neuron | (Butovsky et al., 2014) |
| **miR-676-3p** | | | Astrocyte | (Butovsky et al., 2014) |
| **miR-92a-3p** | Liver | (Guo et al., 2014) | | |
| **miR-92b-3p** | Brain | (Ludwig et al., 2016) | | |
| **miR-93-5p** | Brain | (Guo et al., 2014) | Microglia | (Butovsky et al., 2014) |
| **miR-9-5p** | Brain | (Ludwig et al., 2016, Guo et al., 2014, Landgraf et al., 2007) | Neuron | (Pena et al., 2009) |
| **miR-99a-5p** | Brain | (Lagos-Quintana et al., 2001) | Astrocyte | (Butovsky et al., 2014) |
| **miR-99b-5p** | Brain | (Lagos-Quintana et al., 2001) | Oligodendrocyte | (Butovsky et al., 2014) |

LANDGRAF, P., RUSU, M., SHERIDAN, R., SEWER, A., IOVINO, N., ARAVIN, A., PFEFFER, S., RICE, A., KAMPHORST, A. O., LANDTHALER, M., LIN, C., SOCCI, N. D., HERMIDA, L., FULCI, V., CHIARETTI, S., FOA, R., SCHLIWKA, J., FUCHS, U., NOVOSEL, A., MULLER, R. U., SCHERMER, B., BISSELS, U., INMAN, J., PHAN, Q., CHIEN, M., WEIR, D. B., CHOKSI, R., DE VITA, G., FREZZETTI, D., TROMPETER, H. I., HORNUNG, V., TENG, G., HARTMANN, G., PALKOVITS, M., DI LAURO, R., WERNET, P., MACINO, G., ROGLER, C. E., NAGLE, J. W., JU, J., PAPAVASILIOU, F. N., BENZING, T., LICHTER, P.,

TAM, W., BROWNSTEIN, M. J., BOSIO, A., BORKHARDT, A., RUSSO, J. J., SANDER, C., ZAVOLAN, M. & TUSCHL, T. 2007. A mammalian microRNA expression atlas based on small RNA library sequencing. *Cell,* 129**,** 1401-14.
LAU, P., VERRIER, J. D., NIELSEN, J. A., JOHNSON, K. R., NOTTERPEK, L. & HUDSON, L. D. 2008. Identification of dynamically regulated microRNA and mRNA networks in developing oligodendrocytes. *J Neurosci,* 28**,** 11720-30.
LEE, E. J., BAEK, M., GUSEV, Y., BRACKETT, D. J., NUOVO, G. J. & SCHMITTGEN, T. D. 2008. Systematic evaluation of microRNA processing patterns in tissues, cell lines, and tumors. *RNA,* 14**,** 35-42.
LETZEN, B. S., LIU, C., THAKOR, N. V., GEARHART, J. D., ALL, A. H. & KERR, C. L. 2010. MicroRNA expression profiling of oligodendrocyte differentiation from human embryonic stem cells. *PLoS One,* 5**,** e10480.
LIANG, Y., RIDZON, D., WONG, L. & CHEN, C. 2007. Characterization of microRNA expression profiles in normal human tissues. *BMC Genomics,* 8**,** 166.
LUDWIG, N., LEIDINGER, P., BECKER, K., BACKES, C., FEHLMANN, T., PALLASCH, C., RHEINHEIMER, S., MEDER, B., STAHLER, C., MEESE, E. & KELLER, A. 2016. Distribution of miRNA expression across human tissues. *Nucleic Acids Res,* 44**,** 3865-77.
NATERA-NARANJO, O., ASCHRAFI, A., GIOIO, A. E. & KAPLAN, B. B. 2010. Identification and quantitative analyses of microRNAs located in the distal axons of sympathetic neurons. *RNA,* 16**,** 1516-29.
NELSON, P. T., BALDWIN, D. A., KLOOSTERMAN, W. P., KAUPPINEN, S., PLASTERK, R. H. & MOURELATOS, Z. 2006. RAKE and LNA-ISH reveal microRNA expression and localization in archival human brain. *RNA,* 12**,** 187-91.
PENA, J. T., SOHN-LEE, C., ROUHANIFARD, S. H., LUDWIG, J., HAFNER, M., MIHAILOVIC, A., LIM, C., HOLOCH, D., BERNINGER, P., ZAVOLAN, M. & TUSCHL, T. 2009. miRNA in situ hybridization in formaldehyde and EDC-fixed tissues. *Nat Methods,* 6**,** 139-41.
RAOOF, R., JIMENEZ-MATEOS, E. M., BAUER, S., TACKENBERG, B., ROSENOW, F., LANG, J., ONUGOREN, M. D., HAMER, H., HUCHTEMANN, T., KORTVELYESSY, P., CONNOLLY, N. M. C., PFEIFFER, S., PREHN, J. H. M., FARRELL, M. A., O'BRIEN, D. F., HENSHALL, D. C. & MOONEY, C. 2017. Cerebrospinal fluid microRNAs are potential biomarkers of temporal lobe epilepsy and status epilepticus. *Sci Rep,* 7**,** 3328.
SMRT, R. D., SZULWACH, K. E., PFEIFFER, R. L., LI, X., GUO, W., PATHANIA, M., TENG, Z. Q., LUO, Y., PENG, J., BORDEY, A., JIN, P. & ZHAO, X. 2010. MicroRNA miR-137 regulates neuronal maturation by targeting ubiquitin ligase mind bomb-1. *Stem Cells,* 28**,** 1060-70.
TAN, C. L., PLOTKIN, J. L., VENO, M. T., VON SCHIMMELMANN, M., FEINBERG, P., MANN, S., HANDLER, A., KJEMS, J., SURMEIER, D. J., O'CARROLL, D., GREENGARD, P. & SCHAEFER, A. 2013. MicroRNA-128 governs neuronal excitability and motor behavior in mice. *Science,* 342**,** 1254-8.
THOMPSON, R. C., DEO, M. & TURNER, D. L. 2007. Analysis of microRNA expression by in situ hybridization with RNA oligonucleotide probes. *Methods,* 43**,** 153-61.
WANG, W. X., DANAHER, R. J., MILLER, C. S., BERGER, J. R., NUBIA, V. G., WILFRED, B. S., NELTNER, J. H., NORRIS, C. M. & NELSON, P. T. 2014. Expression of miR-15/107 family microRNAs in human tissues and cultured rat brain cells. *Genomics Proteomics Bioinformatics,* 12**,** 19-30.
